# Reduced taurine transporter expression in lymphoblastoid cell lines from Alzheimer’s disease patients compared with age-matched controls: Therapeutic implications?

**DOI:** 10.1101/2025.03.31.646363

**Authors:** Yuval Gavriel, Irena Voinsky, Hana Klin, Alessio Squassina, David Gurwitz

## Abstract

Taurine is an atypical amino acid that cannot form peptide bonds and thus does not take place in building proteins. Yet, taurine takes part in regulating many cell functions, including cell osmolarity and volume, mitochondrial function, membrane ion channels and neuronal activity, and cell survival. Taurine is synthesized by the liver, and available from consumption of meat and fish, but not plants. It has millimolar concentrations in the brain, skeletal muscle, blood, heart, retina, and other tissues. Taurine is transported from the liver (following synthesis) or the intestine (following consumption) to blood by the taurine transporter, encoded in humans by *SLC6A6*. A recent study reported that blood taurine declines dramatically in aged individuals. Several studies indicated that dietary taurine slows cognitive decline in Alzheimer’s disease (AD) model mice. We therefore measured *SLC6A6* mRNA expression in human lymphoblastoid cell lines (LCLs) from AD patients and age-matched controls and observed 2.8-fold lower expression in AD LCLs (p=0.0005). Additionally, glutathione peroxidase 1 (*GPX1*), a key free-radical scavenging selenoenzyme, had reduced mRNA expression in LCLs from AD patients compared with controls. Our observations suggest that reduced taurine transporter expression may contribute to AD pathogenesis and that dietary taurine might be beneficial for slowing disease progression in early-stage AD. Clinical trials with dietary taurine supplementation of individuals with mild cognitive impairment (MCI) or early-stage AD are required to assess its tentative therapeutic potential.

## 1. Introduction

### 1.1 Taurine: a semi-essential amino acid implicated in healthy aging

Taurine (2-aminoethanesulfonic acid) is a semi-essential micronutrient that regulates various physiological processes. Taurine is an atypical amino acid that contains a sulphonyl group in place of a carboxyl group and therefore cannot make peptide bonds or build proteins. Taurine thus exists primarily as a free amino acid inside cells. Taurine is synthesized from serine in the liver of most mammals, including humans, albeit not always in sufficient amounts. It remains largely unchanged when carried in extracellular body fluids [Huxtable 2000; Bhat et al., 2020].

Since its discovery in 1827, taurine has been studied mostly in the context of protection from reactive oxygen species (ROS) and the regulation of mitochondrial function, ion channels, neuronal activity, protein aggregation, and cell survival [Moenkemann et al., 1999; Surai et al., 2021]. Taurine is also known to be an osmolyte implicated in cell volume regulation. Taurine modulates intracellular free calcium concentration, and even though it is not incorporated into proteins, it is one of the most abundant amino acids in humans, found at millimolar extracellular concentrations in the brain, skeletal muscles, heart, retina, skin, and other tissues [Ripps et al., 2012; Yoshimura et al., 2021]. Taurine is synthesized from serine by three distinct enzymatic steps: serine dehydratase converts serine to 2-aminoacrylate, which is next converted to cysteic acid by 3′-phosphoadenylyl sulfate:2-aminoacrylate C-sulfotransferase. Cysteic acid is next converted to taurine by cysteine sulfinic acid decarboxylase [Sass and Martin., 1972; Martin et al., 1972; Tappaz et al., 1992]. In addition to its synthesis, which takes place mostly in the liver, dietary taurine is available from fish, meat, and dairy products, but not from plants [Huxtable, 2000]. Dietary taurine requires a transporter to be absorbed by the intestinal epithelial cells, moved to the blood’s plasma, and taken up into cells. In humans, the taurine transporter, encoded by *SLC6A6* (Solute Carrier Family 6 Member A6; NCBI Gene ID 6533) is required for taurine export from the liver hepatocytes, its major biosynthesis tissue, for its intestinal uptake from the diet and, eventually, entry into other cell types. The human taurine transporter is ubiquitously and functionally expressed in most tissues and cell types including the intestinal brush border [Anderson et al., 2009], brain [Kaczmarek and Davison, 1972; Yoshimura et al., 2021], retina [Preising et al., 2019], the blood-brain barrier [Tachikawa and Hosoya, 2011], and blood-testis barrier [Kubo et al., 2022]. The particularly high taurine concentrations in the rat heart (approximately 20 mM) are maintained by high expression levels of the taurine transporter [Ito and Murakami, 2024]. In the heart taurine participates in regulating intracellular ion dynamics and calcium handling and has been shown to reduce cellular senescence and increase the lifespan of mice [Singh et al., 2023].

Taurine regulates a variety of processes that are dysregulated during the aging process [Singh et al., 2023]. In their Science article, Singh et al. showed that serum taurine levels decline during aging in several species. In female *C57Bl/6J* wild-type mice, serum taurine levels decreased >3-fold from the age of 4 weeks to 56 weeks. In female monkeys, serum taurine decreased >6-fold from the age of 5 years to 15 years; while in humans, serum taurine levels decreased nearly 5-fold from adulthood to old age [Singh et al., 2023]. These observations are consistent with earlier reports that taurine levels decline with age in rat skin [Yoshimura et al., 2021]. A reversal of the age-associated decline in serum taurine through taurine supplementation from mid-life increased the life span of mice by two months [Singh et al., 2023]. Taurine supplementation reduced cellular senescence and attenuated inflammation, as evident from reduced concentrations of pro-inflammatory cytokines, such as tumor necrosis factor-alpha (TNF-alpha) and interleukin-6 (IL-6) that were increased in middle-aged mice not supplemented with dietary taurine [Yoshimura et al., 2021; Singh et al., 2023]. In young healthy volunteers, taurine reduced the serum pro-inflammatory cytokine IL-6 following a 5 km running exercise [Wang et al., 2022].

### 1.2 Taurine in Alzheimer’s disease

In addition to its implications in healthy aging, taurine is emerging as a beneficial modulator of Alzheimer’s disease (AD), influencing cognitive deterioration, age of onset, and rate of neurodegeneration. Dietary taurine supplementation in drinking water has been shown to improve learning and memory in the adult APP/PS1 mouse model of Alzheimer’s disease [Kim et al., 2014]. Taurine also reduced the neurotoxicity of oligomeric amyloid-β and glutamate receptor agonists in chick retinal neurons (Louzada et al, 2004). It directly bound oligomeric amyloid-β and reduced cognitive decline in Alzheimer’s model mice [Jang et al., 2017]. In the 5xFAD transgenic mouse Alzheimer’s model, which included positron emission tomography imaging studies, taurine supplementation protected mice from reduced brain glutamatergic neurotransmission [Oh et al., 2020]. A recent study used a rat model of hippocampal neuroinflammation, namely, induced by bilateral intra-cerebroventricular (*i*.*cv*.) administration of streptozotocin (STZ), showing that dietary taurine (100 mg/kg orally per day) protected rats against STZ-induced memory impairment [Huf et al., 2024]. Another recent study showed that adding 1% taurine to the drinking water of senescence-accelerated mouse prone 8 (SAMP8) mice reduced the numbers of activated microglia in their hippocampus and cortex, along with reduced phospho-tau and Aβ deposit in these brain regions [Ahmed et al., 2024].

Taurine is the third most abundant amino acid in the brain, after glutamate and glutamine, having a brain concentration of ∼1.2 mM [Griffin and Bradshaw, 2017]. Using immortalized rat brain capillary endothelial cells (TR-BBB13) as an in vitro model of the blood-brain barrier (BBB), sodium-dependent taurine uptake was observed [Kang et al., 2002]. Taurine uptake by these model BBB endothelial cells increased ∼3-fold by TNF-alpha under hypertonic conditions, suggesting that taurine uptake across the BBB is upregulated in response to inflammatory cell damage [Kang et al., 2002]. Together, these studies suggest that dietary taurine might reduce neuronal damage in AD and that its dietary supplementation should be studied by clinical trials in individuals with familial AD risk or those diagnosed with minimal cognitive impairment (MCI). Here, we report our findings on reduced *SLC6A6* expression in lymphoblastoid cell lines (LCLs) generated from blood samples of AD patients compared with non-demented age-matched controls. Our findings should be interpreted with caution until validated in blood or peripheral PBMC samples from AD patients and age-matched controls.

## 2. Patients, Cell Lines, Materials and Methods

### 2.1 Patients and lymphoblastoid cell lines (LCLs)

Alzheimer’s disease (AD) patients and controls were collected at the University Hospital of Cagliari, Italy (Hadar et al., 2016). Demographic data (age at blood collection, age of AD onset, and cognitive scores of the AD patients are presented in **Supplementary Table S1**. LCLs were generated with informed consent from blood mononuclear cells (PBMC) of patients and healthy controls. The cells were grown in parallel under optimal growth conditions before RNA extractions as described in our previous studies (Hadar et al, 2016; Hadar et al, 2018). Tissue-culture reagents were purchased from Sartorius Biological Industries (Beit-Haemek, Israel).

### 2.2 RNA extraction, real-time qPCR, and statistics analysis

RNAs were extracted from LCLs grown in upright T-25 flasks under optimal growth conditions in serum-containing RPMI media at cell densities of 0.5 × 10^6^ to 1 × 10^6^ cells/ml (Voinsky et al., 2024). Cells were centrifuged and then applied for RNA extraction (performed in parallel) using Zymo Research Quick-RNA MiniPrep kit #R1054 (Irvine, CA, USA). RNA quality and yields were determined by NanoDrop Microvolume Spectrophotometer (Thermo Fisher Scientific, MA, USA). RNAs (1 ug from each LCL) were converted to cDNA using qScript cDNA Synthesis Kit (Quanta Bio, Beverly, MA, USA). Reverse transcription was performed using a thermal cycler over three steps (22°C for 5 min, followed by 42 °C for 30 min, and 85 °C for 5 min). Real-time PCR reactions were performed in mixtures containing 10 ng of cDNA, PerfeCTa SYBR^®^ Green FastMix Kit (Quanta Bio, Beverly, MA, USA), and Integrated DNA Technologies, Inc. (Leuven, Belgium) primers. *GUSB* (glucuronidase beta) was used as a reference gene. The cDNAs prepared from the RNAs were applied for real-time qPCR (Hadar et al., 2018, Voinsky et al., 2024) to determine the expression of *SLC6A6* and *GPX1* (glutathione peroxidase 1) in the LCLs from AD patients and age-matched controls. The following SYBR^®^ Green primers were used for the RT-qPCR reactions:

*SLC6A6* forward: TGGTGGAGGTGCGTTTCTC

S*LC6A6* reverse: AGAGGTGTACTGGCCTATGATG

*GPX1* forward: CAGTCGGTGTATGCCTTCTCG

*GPX1* reverse: GAGGGACGCCACATTCTCG

*GUSB* forward: CTGCTGGCTACTACTTGAAGATG

*GUSB* reverse: GAGTTGCTCACAAAGGTCAC

Real-time qPCR data analysis was conducted using GraphPad Prism v.9 (San Diego, CA, USA) software. The Shapiro–Wilk test was used to assess the normality of the data distribution. Differences in continuous variables between the two groups were analyzed using the Mann– Whitney test. Outliers were identified using the ROUT test. A p-value of ≤0.05 was considered statistically significant. Correlations were examined by assessing the normality of data distribution using the Shapiro–Wilk test, followed by a Pearson correlation test (Voinsky et al., 2024).

## 3. Results

Our real-time qPCR measurements indicated reduced mRNA expression of *SLC6A6* (FD=3.28; p<0.001) in LCLs from Alzheimer’s disease patients compared with age-matched controls (**Fig. 1**). Note the large *SLC6A6* expression variation in the control LCLs compared with its smaller variation in AD LCLs (**Fig. 1**). We also observed a positive correlation between lower *SLC6A6* expression levels in LCLs from AD patients and earlier disease onset ages (R=0.449; p=0.0239; **Fig. 2**). However, no correlations were observed between *SLC6A6* expression in LCLs from AD patients and their cognitive scores (**Supplementary Fig. S1**).

**Figure 1:**
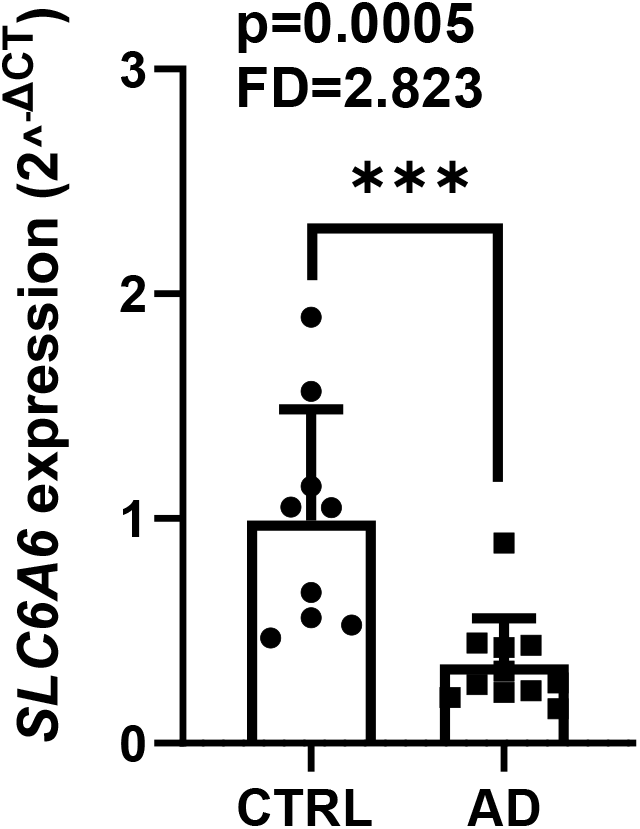
Reduced *SLC6A6* expression in lymphoblastoid cell lines (LCLs) from Alzheimer’s disease (AD) patients vs. age-matched healthy controls. RNAs were extracted from LCLs of AD patients and age-matched healthy controls. Average ages (at the time of blood sample collection for LCL generation) were closely similar for the two groups: 72.5 and 71.7 years for the AD patients and the controls, respectively (Supplementary Table S1). Following cDNA preparation from the extracted RNAs, real-time qPCR experiments were performed (see Methods). As shown, the AD LCLs expressed 2.8-fold lower *SLC6A6* mRNA levels compared with LCLs from the age-matched controls (p=0.0005).

**Figure 2:**
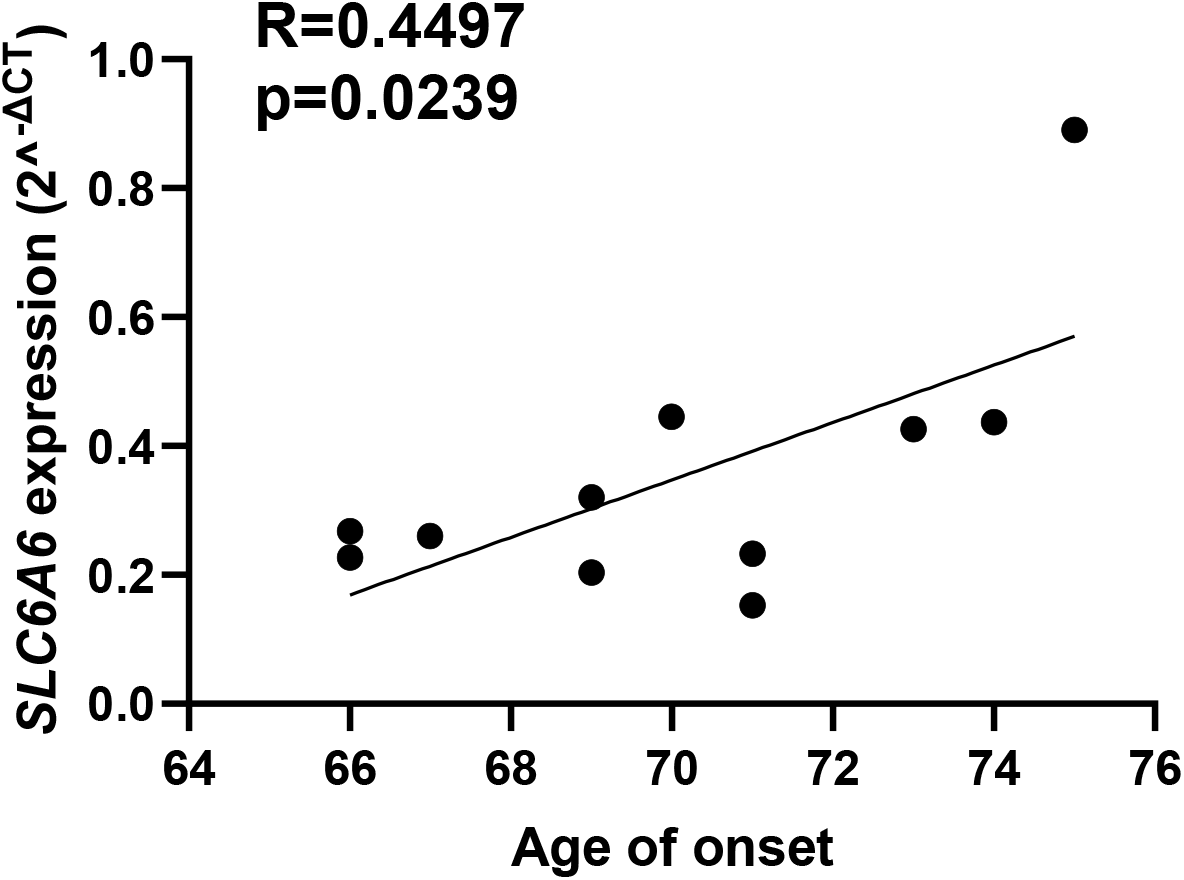
*SLC6A6* expression levels in lymphoblastoid cell lines from Alzheimer’s disease patients positively correlate with their AD age of onset. *SLC6A6* mRNA expression levels in LCLs from AD patients (Figure 1) were plotted against the age of AD onset. A positive correlation was found between these parameters. See Methods for further details.

We also measured glutathione peroxidase 1 (*GPX1*) mRNA in the AD and control LCLs, as we recently reported this gene, coding for a selenoprotein taking part in reactive oxygen species (ROS) inactivation, with decreased expression in LCLs from older individuals (Voinsky et al., 2024). *GPX1* codes for a selenoenzyme that plays a key role in protecting cells against reactive oxygen species free radicals [Jablonska et al., 2009]. We observed lower *GPX1* mRNA expression in LCLs from AD patients compared with age-matched healthy controls (FD=1.76; p=0.034; **Fig. 3**). As both *SLC6A6* and *GPX1* showed decreased mRNA expression in LCLs from AD patients compared with age-matched controls, we examined the correlation between their expression levels. Remarkably, we found a positive correlation between the mRNA expression of *SLC6A6* and *GPX1* in the control LCLs (p=0.0019; **Fig. 4a**); while no correlations were observed between the expression levels of these two genes in LCLs from AD patients (**Fig. 4b**).

**Figure 3:**
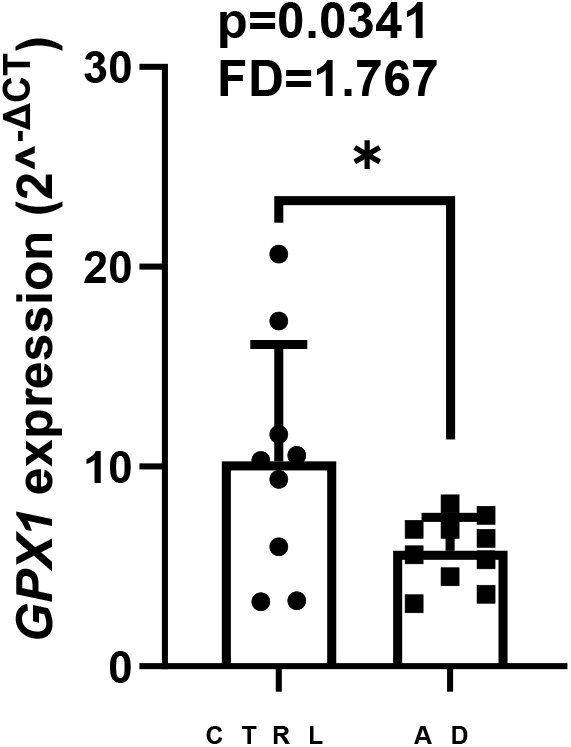
Reduced *GPX1* expression in lymphoblastoid cell lines from Alzheimer’s disease patients compared with age-matched controls. The same RNAs extracted from the AD and control LCLs that were applied for Fig. 1 were used. See Fig. 1 for further details.

**Figure 4:**
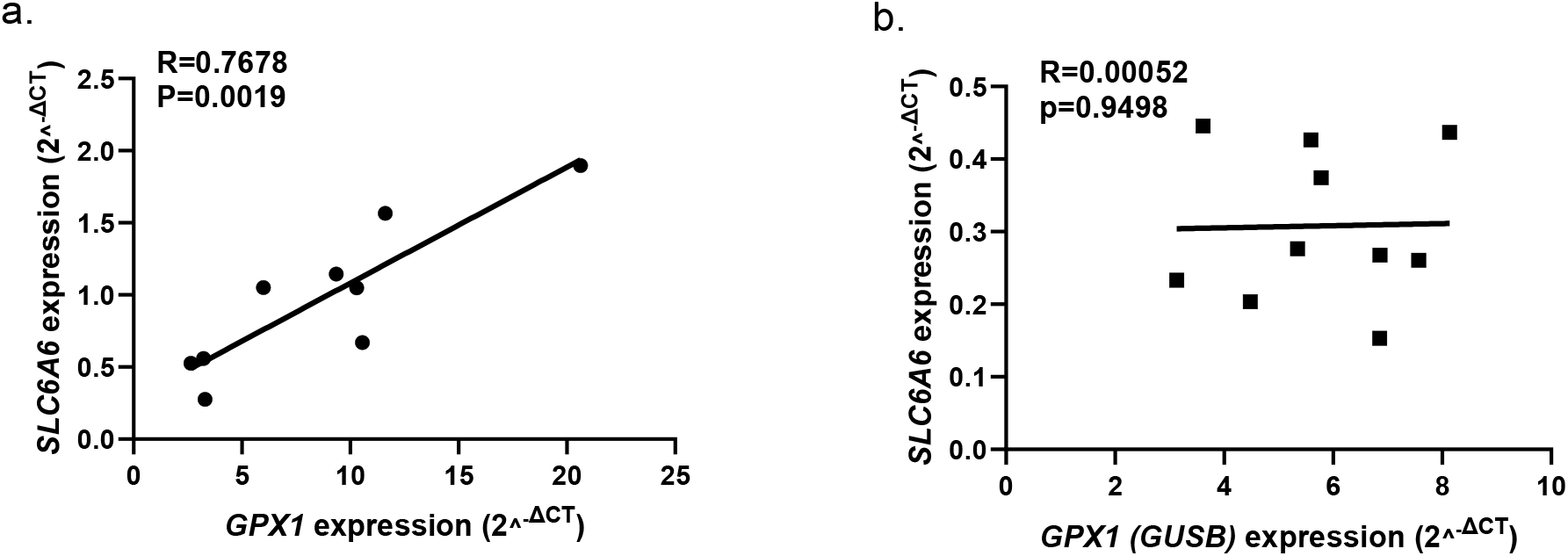
Correlation between *SLC6A6* and *GPX1* mRNA expression in control and AD lymphoblastoid cell lines. (**a**) Control LCLs show a strong correlation between the expression of *SLC6A6* and *GPX1* mRNA levels. **(b**) AD LCLs showed no such correlation; one outlier was removed. Expression levels were taken from Fig. 1 and Fig. 3 for *SLC6A6* and *GPX1*, respectively.

## 4. Discussion

### 4.1 Reduced expression of taurine transporter (*SLC6A6*) in AD LCLs

The taurine transporter, coded in humans by *SLC6A6*, is essential for the export of taurine from the liver, where it is synthesized from serine, and for taurine adsorption from the diet by gut epithelial cells [Huxtable, 2000; Singh et al., 2023; Ito and Murakami, 2024]. Thus, our findings on reduced *SLC6A6* mRNA expression in LCLs from AD patients compared with age-matched control LCLs (**Fig. 1**) agree with studies proposing that dietary taurine is beneficial for slowing cognitive decline in AD rodent models [Kim et al., 2014; Jang et al., 2017; Oh et al., 2020; Huf et al., 2024]. However, these suggestions were based on dietary taurine supplementation in model AD rodents, while our current findings with LCLs from AD patients compared with age-matched controls provide the first clue for reduced taurine availability in AD patients. However, proof of a direct link between lower taurine transporter expression and reduced taurine availability in AD patient tissues will require measurements of taurine levels in AD patients’ blood samples, PBMCs, or postmortem brain tissues.

Serum vitamin B12 deficiency was associated with cognitive functioning in elderly individuals [Robins Wahlin et al., 2001; Brito et al., 2017]. Lower serum vitamin B12 levels were observed in AD patients [Ikeda et al., 1990] and correlated with their Mini Mental State Examination (MMSE) scores [Levitt and Karlinsky, 1992]. It is therefore notable that vitamin B12 is essential for taurine synthesis by the liver [Roman-Garcia et al., 2014], and that low serum B12 was associated with low serum taurine in a metabolomics study of a human cohort [Baghel et al., 2024].

We observed a positive correlation between *SLC6A6* expression in LCLs from AD patients and the age of onset of their disease (**Fig. 2**), while no correlation was found between *SLC6A6* expression and cognitive scores in AD patients (Supplementary **Fig. S1**). The correlation between AD age of onset and the disease progression rate remains controversial. Earlier studies reported a lack of such correlations (Huff et al., 1987; Haupt et al., 1993; Bowler et al., 1998). While later studies observed such a correlation and suggested that an earlier age of AD onset could be prognostic for a faster rate of disease progression (Sinforiani et al., 2019; Day et al., 2022).

### 4.2 Reduced expression of glutathione peroxidase 1 (*GPX1*) in AD LCLs

We recently reported that three selenoproteins, including *GPX1*, exhibited reduced mRNA expression in LCLs from older women (Voinsky et al., 2024). In addition, reduced glutathione peroxidase activity was observed in blood samples from AD patients compared to age-matched controls, while its levels were not correlated with the patient’s cognitive scores (Garlet et al., 2019). Our observations of lower expression of *GPX1* in the AD LCLs compared with their control LCLs (**Fig. 3**) agree with the central concept that oxidative stress is a driver of aging and AD (Bai et al., 2022), and with suggestions that antioxidants and anti-inflammatory compounds may slow AD progression, albeit other free radicals may contribute to neuronal cell death in AD (Mani et al., 2023).

In this context, our findings on a positive correlation between *SLC6A6* and *GPX1* expression in control LCLs (**Fig. 4a**), but not in AD LCLs (**Fig. 4b**) are puzzling. They suggest that shared mechanism(s) may regulate the transcription of both genes and may become dysfunctional in AD. A shared miRNA is unlikely to be implicated: miRBase (www.mirbase.org), an archive for microRNA sequences and their target annotations, does not include shared and evolutionally preserved human miRNA predicted to bind to the 3’UTR regions of both *SLC6A6* and *GPX1*. Notably, metabolic network analysis of post-mortem brains from AD patients and non-demented controls that focused on alternative bile acid synthesis pathways suggested that bile acid synthesis, taurine transport, and cholesterol metabolic pathways differed in AD and controls [Baloni et al., 2020]. Future studies should explore, among other options, the possibility that differential promoter methylation, or another epigenomic mechanism, is implicated in these perplexing observations.

### 4.3 Implications for neurodegenerative disorders

Taurine has been shown to promote autophagy in matured 3T3-L1 mouse adipocytes [Kaneko et al., 2018], while taurine deficiency was associated with dysfunctional autophagy and ubiquitination in the heart of mice with deleted taurine transporter gene [Jong et al., 2015]. Autophagy allows the lysosome-dependent degradation and recycling of dysfunctional proteins to allow cellular renovation [Mizushima and Komatsu, 2011]. It also plays a key role in removing aggregated or misfolded proteins, including aggregated amyloid-beta and phosphorylated tau that accumulate in AD brain tissues [Lipinski et al., 2010; Tian et al., 2013]. Dysfunctional autophagy is also implicated in Parkinson’s and Gaucher diseases [Nechushtai et al., 2023; Hull et al., 2024], as well as in diabetes [Goldman et al., 2010; Chen et al., 2011]. Thus, the reduced *SLC6A6* expression levels observed in AD LCLs may affect the reduced autophagy biomarkers reported in sera from Alzheimer’s patients (Castellazzi et al., 2019).

Animal studies reported that dietary taurine supplementation may protect from additional neurological diseases or disorders besides AD. Taurine improved cognitive impairment and neuronal cell death in model Parkinson’s disease (PD) mice or rats by inhibiting microglial activation [Wang et al., 2021] and alpha-synuclein aggregation [Onuelu et al., 2025]. Lower blood taurine was reported in levodopa-treated PD patients [Zhang et al., 2016]. Lastly, taurine supplementation also protected mice from axonal degeneration following spinal cord injury [Sobrido-Cameán et al., 2020].

Among the inspiring studies on the biology of taurine and its transporter is a study showing that immune exhaustion in CD8+ T lymphocytes, and thereby immune escape of aggressive tumors, is driven by cancer-related taurine consumption [Cao et al, 2024]. These authors showed that cancers express elevated levels of the taurine transporter, thereby competing with T cells on the limited supply of taurine. In other words, SLC6A6-mediated taurine deficiency encourages immune evasion of tumor cells. Dietary taurine supplementation bolsters the exhausted CD8+ T cells and allows them to overcome the immune evasion of the tumor cells. If proven correct in clinical trials, dietary taurine could improve the efficacy of cancer therapies.

It remains unclear how diminished taurine availability contributes to neuronal damage in AD, PD, spinal cord injury, and other neurodegenerative disorders, or to tumor cells immune evasion. The paucity of human studies and, ultimately, clinical trials of taurine supplementation, may reflect a lack of financial incentives for the pharma sector to invest in clinical trials with dietary taurine – a cheap natural compound that cannot be patented. Ultimately, such clinical trials may require funding by public organizations or grant agencies. A plausible path for encouraging the private sector interest in taurine or taurine mimetics would require the development and patenting of cell-permeable taurine-liposome or taurine precursor formulations converted to taurine by intracellular enzyme(s), thereby overcoming conditions of reduced cellular taurine transporter expression. Indeed, out of 82 clinical trials with taurine found in clinicaltrials.gov (search performed on March 30, 2025), none was directed at AD, PD, MCI, or spinal cord injury. Rather, there were seven trials related to cardiovascular diseases, six for diabetes, and five for the generic term “aging”. The latter include a recruiting clinical trial titled “Effects of daily taurine intake for 6 months on biological age and body metabolism indicators as well as physical fitness in 55-75-year-old women and men” (NCT06613542) sponsored by the Technical University of Munich, Germany. Similar trials with AD and MCI patients seem warranted.

A notable limitation of our study is using n vitro measurements in LCLs, rather than MCI or AD biosamples such as blood, PBMC, or buccal swabs. LCLs are immortalized human B lymphocytes that represent a small fraction of PBMCs. While B lymphocytes differ from CNS neuronal or glial cells, they were useful for research on CNS disorders, including AD [Hadar et al., 2016; Hadar et al., 2018] and the mode of action of antidepressants [Oved et al., 2013]. LCL repositories serve as a useful resource for generating induced pluripotent stem cells (iPSCs) [Gurwitz, 2016; Gurwitz and Steeg, 2014]. Importantly, LCLs take up labeled taurine added to their growth medium [Tallan et al., 1983, Lin et al., 1988], suggesting that taurine may affect antibody production, a capacity known to decline in old age [Frasca and Blomberg, 2016].

## 5. Conclusions and Future Outlook

To our knowledge, this is the first study to report reduced expression of the taurine transporter,

*SLC6A6*, in cells derived from AD patients compared with age-matched controls. Future studies and ultimately clinical trials should assess whether long-term dietary taurine supplementation or upregulation of *SLC6A6* expression could slow disease progression in early-stage AD or MCI patients without adverse effects. Meanwhile, our findings on reduced taurine transporter expression in AD-derived cells should be interpreted with caution.

## Acknowledgments

The authors are grateful for the individuals who consented to donate blood samples for the generation of LCLs, which made this study possible. This study is part of the M.Sc. project of Hana Klin at the Graduate School of the Faculty of Medical and Health Sciences, Tel Aviv University, Israel.

## Data Availability Statement

The data generated for the current study are available from the corresponding author on reasonable request.

## Conflicts of Interest

The authors declare no competing interests that might influence the results and/or discussion reported in this article.

## Notes

### Competing Interest Statement

The authors have declared no competing interest.

