## Supplemental Table 1 and Figure S1 for "Reduced taurine transporter expression in lymphoblastoid cell lines from Alzheimer’s disease patients compared with age-matched controls: Therapeutic implications?"

**Supplementary Table S1**: **Demographics of Alzheimer’s disease (AD) LCL donors**.

Average ages at time of blood withdrawal for LCL preparation were 72.5 and 71.7 years for the AD patients and the controls, respectively. All AD patients were female, similarly to the donors applied for our earlier study (Hadar et al., 2016). ^*^Age (years) was at time of blood withdrawal for LCL generation. MMSE and ADAS scores were taken on the day of blood withdrawal. Four of the control LCLs (codes starting with 5) were from Israeli LCL collection for achieving closely matching average age of controls to the AD patients (see: <https://www.nlgip-yoran.sites.tau.ac.il/>).

Alzheimer’s disease patients

LCL	Age^*^	Age at onset	MMSE	ADAS

1110	75	73	24	11.9

1130	76	75	20.7	10.6

1141	75	73	22	23

1197	71	69	19.7	10.6

1145	68	66	NA	NA

1135	75	70	22.3	17.2

1090	73	71	21.4	14.9

1118	68	67	21.4	19.7

1100	70	66	14.3	23.2

1121	75	74	19.7	15.6

1173	71	69	19.3	23.3

1229	73	71	20.4	20.3

Age-matched controls

1060C	79

1377C	82

1378C	70

1379C	74

1380C	74

1446C	63

5096C	74

5097C	77

5199C	59

5922C	65

**Supplementary Fig. S1. Lack of correlations between** ***SLC6A6*** **mRNA** **expression levels in LCLs from Alzheimer’s disease patients and their cognitive scores at times of blood sample collection.** (**a**) *SLC6A6* mRNA expression vs. ADAS scores; (**b**) *SLC6A6* mRNA expression vs. MMSE scores. Cognitive scores were measured on the day of blood withdrawal for LCL generation. As shown, there were no correlations between either score and *SLC6A6* mRNA expression in LCLs derived from AD blood samples. See Methods for further details.

**a.	b.**

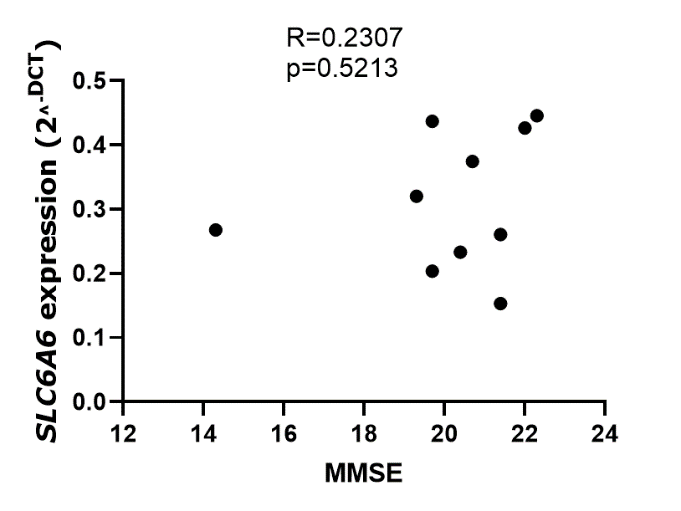

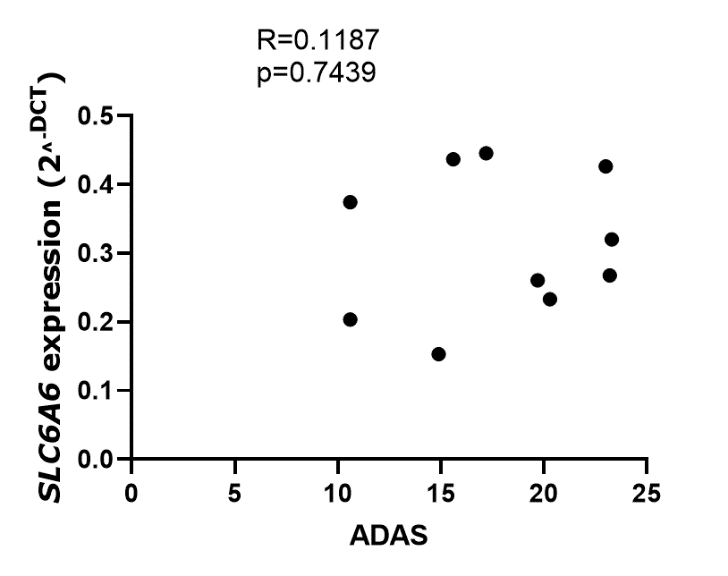
